# *Listeria monocytogenes* requires DHNA-dependent intracellular redox homeostasis facilitated by Ndh2 for survival and virulence

**DOI:** 10.1101/2023.01.13.524026

**Authors:** Hans B. Smith, Kijeong Lee, David M. Stevenson, Daniel Amador-Noguez, John-Demian Sauer

## Abstract

*Listeria monocytogenes* is a remarkably well-adapted facultative intracellular pathogen that can thrive in a wide range of ecological niches. *L. monocytogenes* maximizes its ability to generate energy from diverse carbon sources using a respiro-fermentative metabolism that can function under both aerobic and anaerobic conditions. Cellular respiration maintains redox homeostasis by regenerating NAD^+^ while also generating a proton motive force (PMF). The end products of the menaquinone (MK) biosynthesis pathway are essential to drive both aerobic and anaerobic cellular respiration. We previously demonstrated that intermediates in the MK biosynthesis pathway, notably 1,4-dihydroxy-2-naphthoate (DHNA), are required for the survival and virulence of *L. monocytogenes* independent of their role in respiration. Furthermore, we found that restoration of NAD^+^/NADH ratio through expression of water-forming NADH oxidase (NOX) could rescue phenotypes associated with DHNA deficiency. Here we extend these findings to demonstrate that endogenous production or direct supplementation of DHNA restored both the cellular redox homeostasis and metabolic output of fermentation in *L. monocytogenes*. Further, exogenous supplementation of DHNA rescues the *in vitro* growth and *ex vivo* virulence of *L. monocytogenes* DHNA-deficient mutants. Finally, we demonstrate that exogenous DHNA restores redox balance in *L. monocytogenes* specifically through the recently annotated NADH dehydrogenase Ndh2, independent of the extracellular electron transport (EET) pathway. These data suggest that the production of DHNA may represent an additional layer of metabolic adaptability by *L. monocytogenes* to drive energy metabolism in the absence of respiration-favorable conditions.

## INTRODUCTION

*Listeria monocytogenes* is a Gram-positive, facultative intracellular pathogen that is exceptionally well-adapted to survive and replicate in the restrictive mammalian host cytosol (1-3). Bacteria that lack the specific adaptations required to survive or replicate in the host niche are effectively cleared (4-7), often by triggering host defense mechanisms comprised of innate immune pathways (8-13). *L. monocytogenes* utilizes its internalin proteins to facilitate invasion into the host cell where it becomes captured in a phagosome (14, 15). The pore-forming cytolysin listeriolysin O (LLO) then facilitates escape from the phagosome into the cytosol (14, 16), where *L. monocytogenes* can utilize ActA to mediate actin-based motility by hijacking the host’s actin machinery (17-20). Using this motility, *L. monocytogenes* moves into adjacent cells where they again invade the cytosol by expressing LLO and two phospholipase Cs, PlcA and PlcB, enabling it to restart its life cycle (14, 21).

*L. monocytogenes* can also thrive in a diverse range of ecological niches that contain highly variable pools of fermentable and non-fermentable carbon sources (2, 22). *L. monocytogenes* employs both fermentative and respiratory metabolic mechanisms to maximize its energy output from scavenged nutrients (22, 23). In contrast to canonical respiratory organisms however, *L. monocytogenes* contains an incomplete tricarboxylic acid (TCA) cycle and is therefore unable to fully oxidize its carbon substrates (24). Accordingly, *L. monocytogenes* utilizes a respiro-fermentative metabolism characterized by glycolysis-derived pyruvate that is funneled into the fermentative production of acetate, generating ATP through substrate-level phosphorylation (SLP) via the activity of acetate kinase (24, 34). During the respiro-fermentative process, the activity of *L. monocytogenes*’ respiratory electron transport chain (ETC) enables it to regenerate NAD^+^, without having to rely upon lactate dehydrogenase, while also producing a functional proton motive force (PMF) (22, 24, 34). Further lending to its diverse metabolic adaptability, *L. monocytogenes* possesses two distinct respiratory ETCs that allow it to respire both aerobically and anaerobically (25). The aerobic ETC in *L. monocytogenes* mediates electron transfer from a type II NADH dehydrogenase, Ndh1, to a membrane-bound menaquinone (MK) and subsequently to terminal cytochrome oxidases QoxAB (aa3) or CydAB (bd) for final transfer to O_2_ (26, 27). In contrast, the recently annotated anaerobic respiratory pathway in *L. monocytogenes* uses a flavin-based ETC to drive extracellular electron transfer (EET) to extracytosolic acceptors such as fumarate or ferric ion using a novel NADH dehydrogenase (Ndh2) and an alternative demethylmenaquinone (DMK) intermediate (25, 28). Both of the respiratory ETC in *L. monocytogenes* rely upon the MK biosynthesis pathway to generate their respective quinone electron acceptors, with the biosynthetic intermediate 1,4-dihydroxy-2-naphthoate (DHNA) functioning as a mutual branching point (**Fig. S1**) (25).

The requirement for *L. monocytogenes* to perform cellular respiration during infection has been well documented (29-32). However, understanding the specific contributions of maintaining cellular redox homeostasis via NAD^+^ regeneration versus the production of a functional PMF to achieve virulence has remained elusive. Further complicating our ability to dissect the specific contributions that cellular respiration may have during infection, the MK intermediates DHNA-CoA and DHNA have recently been reported to be required for the survival and virulence of *L. monocytogenes* independent of MK synthesis and aerobic respiration (29, 31, 32). Importantly, although it was observed that the supplementation of exogenous DHNA could rescue the *in vitro* growth of a DHNA-deficient *L. monocytogenes* mutant, this rescue did not coincide with the restoration of its PMF (31). Therefore, although DHNA-deficient strains of *L. monocytogenes* possess the downstream enzymes to produce MK or DMK, these data suggest that exogenous DHNA is not utilized to promote either aerobic or anaerobic cellular respiration. Recent work from Rivera-Lugo *et al*. sought to dissect the relative importance of maintaining redox homeostasis versus PMF generation for the pathogenesis of *L. monocytogenes* using a water-forming NADH oxidase (NOX) that specifically regenerates NAD^+^ independent of respiration and PMF function (34). Through the heterologous expression of NOX in respiration-deficient strains of *L. monocytogenes*, it was concluded that the regeneration of NAD^+^ represents a major role for cellular respiration during pathogenesis.

The studies presented here sought to define the respiration-independent mechanisms of DHNA utilization to promote the survival and virulence of *L. monocytogenes*. Consistent with observations from Rivera-Lugo *et. al*, in the absence of respiration, the *ex vivo* and *in vivo* virulence defects associated with DHNA-deficiency were a result of impaired redox homeostasis which could be rescued upon ectopic NOX expression. Similarly, exogenous DHNA supplementation rescues the *in vitro* and *ex vivo* growth and cytosolic survival of DHNA-deficient mutants. Indeed, DHNA-dependent rescue by direct supplementation resulted in a restored cellular redox homeostasis with a concurrent shift of fermentative flux from lactate production to acetate in *L. monocytogenes*, independent of respiration. We further go on to show that the recently annotated anaerobic-specific Ndh2 is essential for DHNA-deficient *L. monocytogenes* mutants to utilize exogenous DHNA for growth in defined medium, independent of its canonical role in EET, suggesting that Ndh2 is the NADH dehydrogenase specifically required for the restoration of redox homeostasis via DHNA. Taken together, these data suggest that the endogenous production of DHNA can be utilized by *L. monocytogenes* to restore both its intracellular redox homeostasis and fermentative metabolic flux through an undefined mechanism requiring Ndh2.

## RESULTS

### Redox homeostasis via NOX shifts fermentative output and rescues *in vitro* growth of DHNA-deficient *L. monocytogenes*

Two main outcomes of cellular respiration include 1) maintaining intracellular redox homeostasis by regenerating NAD^+^ from NADH and 2) the generation of a PMF to drive oxidative phosphorylation and various other aspects of bacterial physiology. A recent study employed a water-forming NADH oxidase (NOX) expression system in *L. monocytogenes* to dissect the relative importance of cellular respiration in maintaining redox homeostasis versus PMF generation (34). We had previously demonstrated that *L. monocytogenes* mutants lacking the key MK biosynthetic intermediate DHNA were attenuated, in part, independent of loss of respiration (29, 31, 32). We hypothesized that restoration of NAD^+^ pools might rescue these virulence defects similar to the rescue observed for mutants lacking components of the respiratory chains (34). To test this hypothesis, we assessed NAD^+^/NADH levels in Δ*menB*, Δ*menI* and Δ*menA* mutants +/-expression of NOX *in trans*. The inability to generate endogenous DHNA by the Δ*menB* mutant results in a severely diminished redox homeostasis as measured by the ratio of oxidized NAD^+^ to reduced NADH. This imbalance was significantly restored by ectopic expression of NOX to a level similar to the Δ*menA* mutant (**Fig. 1A**). The Δ*menI* mutant, which can generate DHNA-CoA, displays an intermediate phenotype between Δ*menB* and Δ*menA* levels, which is similarly rescued upon NOX expression (**Fig. 1A**), consistent with possible respiration independent roles for DHNA in NAD^+^/NADH redox balancing.

**Figure 1.**
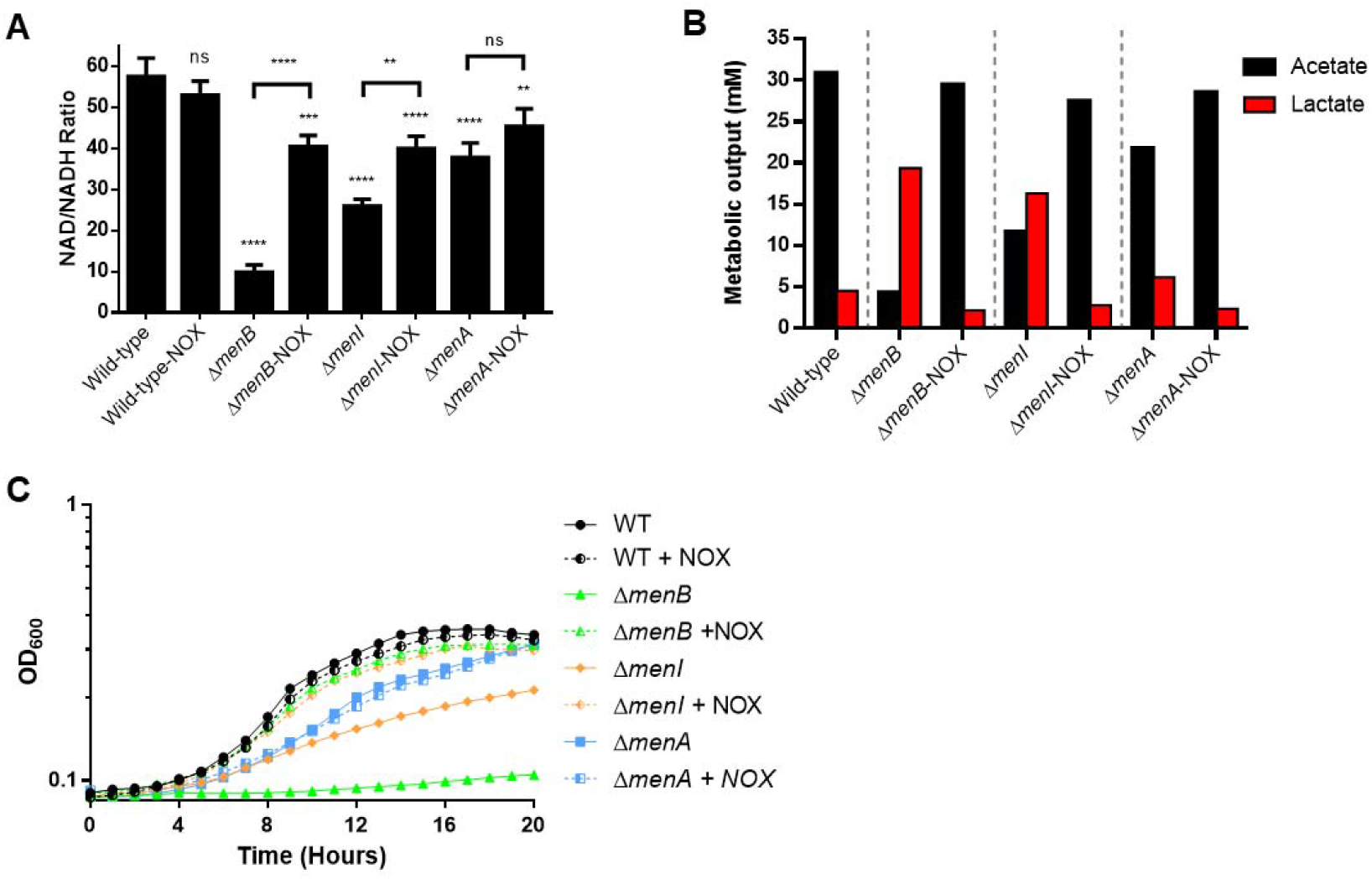
Redox homeostasis via NOX shifts fermentative output and rescues *in vitro* growth of DHNA-deficient *L. monocytogenes*. (A) NAD^+^/NADH ratios of indicated *L. monocytogenes* strains +/-NOX plasmid complementation grown aerobically at 37°C in defined medium to mid-logarithmic phase (OD_600_ 0.4-0.6). Δ*menB* mutant fails to grow in defined medium, thus these culture samples were spiked with 2 × 10^8^ total CFU from an overnight BHI culture during experimental setup. (B) HPLC quantification of fermentation products (Lactate and Acetate) produced and secreted by indicated *L. monocytogenes* strains +/-NOX plasmid complementation grown in BHI media aerobically at 37°C to stationary phase. (C) *L. monocytogenes* strains +/-NOX plasmid complementation were grown in defined medium at 37°C. OD_600_ was monitored for 20 hours. Data are representative of three (A, C) or two (B) independent experiments. ns, not significant; WT, wild-type

*L. monocytogenes* employs a respiro-fermentative metabolism due to an incomplete TCA cycle, characterized by the funneling of pyruvate towards the fermentative production of acetate (23, 24). Respiration-deficient mutants of *L. monocytogenes* are impaired in their ability to maintain cellular redox homeostasis and as a result nearly exclusively produce lactate rather than acetate as a metabolic byproduct (34). To test whether impaired redox homeostasis due to DHNA-deficiency would similarly result in the predominant production of lactate, we analyzed fermentation byproducts in bacterial supernatants using high-performance liquid chromatography (HPLC). As expected, wild-type *L. monocytogenes* predominantly generated acetate whereas DHNA-deficient Δ*menB* had a drastic shift to lactate production (**Fig. 1B**). Heterologous NOX expression rescued Δ*menB* acetate production back to wild-type levels, consistent with restored redox homeostasis driving acetate production to generate ATP (**Fig. 1B**). Consistent with the results seen in our NAD^+^/NADH experiments, the Δ*menI* mutant displayed an intermediate phenotype by producing similar levels of acetate and lactate, which was also fully restored to wild-type upon NOX expression (**Fig. 1B**). The Δ*menA* mutant produced slightly more lactate and less acetate when compared to wild-type, likely attributed to the difference in redox homeostasis observed previously (**Fig. 1A, B**).

Finally, we have previously shown that the production of DHNA is critical for *L. monocytogenes in vitro* growth in chemically defined medium (29, 31, 32). To test whether restoration of redox homeostasis can rescue this growth defect, we’ve assayed for *in vitro* growth of the above mutants complemented with NOX in defined medium. As expected, Δ*menB* showed the largest growth defect followed by Δ*menI*, and both mutants showed wild-type level growth upon NOX complementation (**Fig. 1C**). Together, these data suggest that metabolic defects associated with DHNA deficiency in *L. monocytogenes* are due to NAD^+^/NADH redox imbalances and that restoration of this balance can rescue Δ*menB* mutant growth and carbon metabolism in *L. monocytogenes*.

### Restoration of redox homeostasis rescues virulence defects associated with DHNA-deficiency

Based on the restoration of *in vitro* growth of Δ*menB* mutants via expression of NOX, we hypothesized that restoration of NAD^+^ pools would similarly rescue virulence defects of DHNA-deficient mutants. DHNA-deficient mutants are susceptible to cytosolic killing in the macrophage cytosol, therefore we assessed cytosolic survival of Δ*menB*, Δ*menI*, and Δ*menA* with or without expression of NOX *in trans* (29, 35). As hypothesized, Δ*menB* and Δ*menI* displayed increased cytosolic killing and NOX expression rescued their survival in the macrophage cytosol (**Fig. 2A**). Rescue by NOX expression was specific to mutants with disrupted NAD^+^/NADH redox homeostasis as NOX expression was unable to rescue cytosolic survival of a Δ*glmR* mutant susceptible to cytosolic killing due to cell wall defects (**Fig 2A**) (33, 35, 36). Consistent with NAD^+^ pool restoration supporting cytosolic survival, Δ*menB* mutant replication in the macrophage cytosol was also rescued upon expression of NOX *in trans* (**Fig. 2B**).

**Figure 2.**
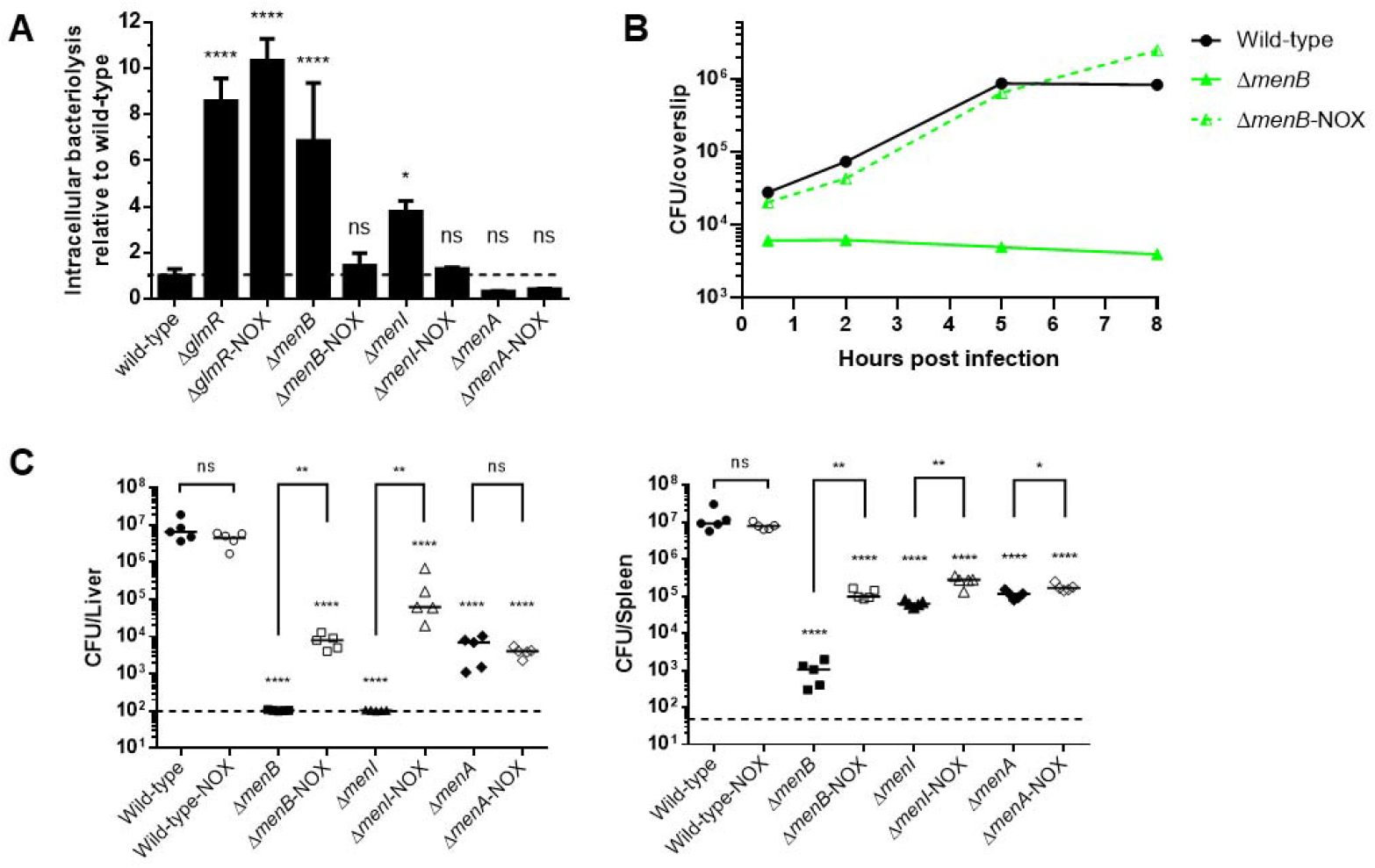
Restoration of redox homeostasis rescues virulence defects associated with DHNA-deficiency. (A) Indicated *L. monocytogenes* strains (MOI of 10) +/-NOX plasmid complementation were tested for cytosolic survival in immortalized IFNAR^-/-^ bone marrow-derived macrophages (BMDM) over a 6 hr infection. Data are normalized to wild-type levels of bacteriolysis and presented as the standard deviation of the means from three independent experiments. (B) Intracellular growth of wild-type, Δ*menB*, or Δ*menB*-NOX was determined in BMDMs following infection at an MOI of 0.2. Growth curves are representative of at least three independent experiments. Error bars represent the standard deviation of the means of technical triplicates within the representative experiment. (C) Bacterial burdens from the spleen and liver were enumerated at 48 hr post-intravenous infection with 1 × 10^5^ total CFU of indicated *L. monocytogenes* strains +/-NOX plasmid complementation. Data are representative of results from two independent experiments. Horizontal bars represent the limits of detection and the bars associated with the individual strains represents the mean of the group. ns, not significant.

Finally, we had previously demonstrated that DHNA-deficient mutants are more attenuated *in vivo* than respiration-deficient mutants, suggesting that DHNA contributes to virulence in a respiration independent manner (29, 31, 32). To determine if the respiration independent function of DHNA during *in vivo* infection is due to NAD^+^/NADH homeostasis defects, we assessed virulence of Δ*menB*, Δ*menI*, and Δ*menA* mutant *L. monocytogenes* with and without expression of NOX *in trans*. Ectopic NOX expression rescued the *in vivo* burden of Δ*menB* mutants by ∼100-fold in the spleen and liver (**Fig. 2C**) and a similar rescue for Δ*menI* mutants in the liver following NOX expression is also observed (**Fig. 2C**). Interestingly, there was little to no change in the *in vivo* virulence of Δ*menA* upon the introduction of NOX (**Fig 2C**). This is in agreeance with our previous results that showed both redox homeostasis and acetate production of the Δ*menA* mutant was also not significantly altered upon NOX expression (**Fig 1A, B**). Taken together, these data suggest that in *L. monocytogenes* maintaining cellular redox homeostasis in the absence of DHNA is sufficient to promote survival and virulence both *ex vivo* and *in vivo*.

### DHNA production or supplementation promotes similar effects to NOX complementation in *L. monocytogenes*

We have previously demonstrated that exogenous addition of either purified DHNA or culture supernatant from DHNA sufficient strains of *L. monocytogenes* could rescue the *in vitro* growth of DHNA-deficient *L. monocytogenes* in defined media (31), suggesting that *L. monocytogenes*, like other bacteria including *Propionibacterium* spp. and *Lactobacillus* spp., may secrete DHNA (43, 45). To test the hypothesis that *L. monocytogenes* secretes DHNA, we assayed culture supernatants for DHNA via mass spectrometry. As hypothesized, wild-type *L. monocytogenes* contained abundant levels of DHNA, while Δ*menB* mutants contained no detectable extracellular DHNA (**Fig. S2)**. Given that exogenous DHNA could rescue the *in vitro* growth of DHNA-deficient *L. monocytogenes* mutants and that DHNA-deficient mutants could similarly be rescued by NAD^+^ regeneration through NOX expression, we hypothesized that exogenous DHNA could act to restore NAD^+^ levels in Δ*menB* mutants. To test this hypothesis, we measured cellular NAD^+^/NADH with or without DHNA supplementation. Consistent with the results observed with NOX expression, the exogenous supplementation of DHNA rescued redox homeostasis of Δ*menB* mutants to levels similar to those seen with Δ*menA* mutants, suggesting that exogenous DHNA might be utilized in a similar fashion to DHNA produced endogenously (**Fig. 3A**). Consistent with DHNA supplementation of Δ*menB* rescuing cellular redox homeostasis, exogenous DHNA also shifted the metabolic flux of Δ*menB* back towards acetate production, similar to Δ*menA* levels (**Fig. 3B**). Importantly, we had previously demonstrated that exogenous DHNA does not restore respiration and membrane potential (31). Taken together, these data suggest that DHNA, independent of its role in respiration, restores cellular redox homeostasis, subsequently shifting the fermentative output from lactate back towards acetate that likely drives ATP production through acetate kinase (24, 34).

**Figure 3.**
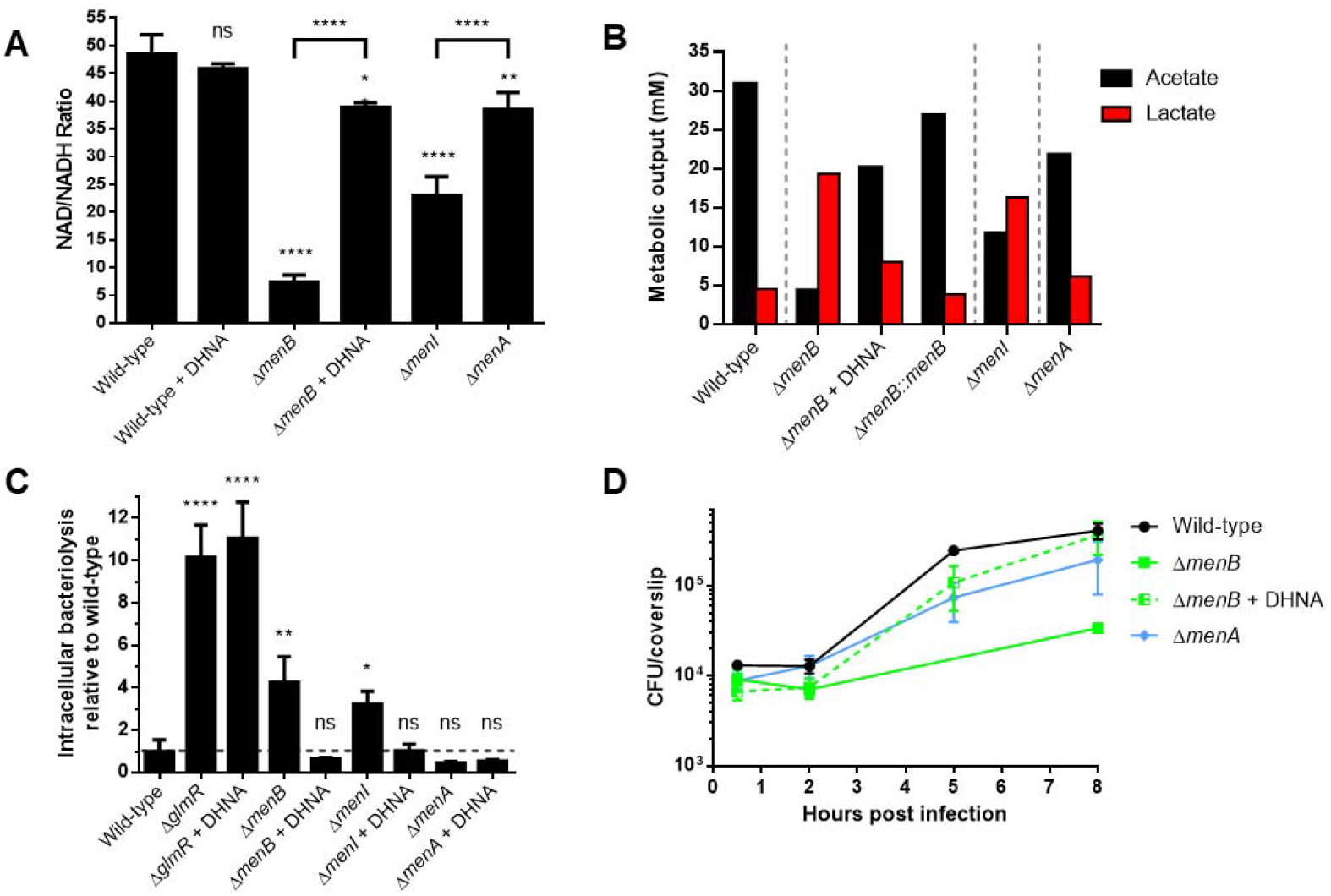
DHNA production or supplementation promotes similar effects to NOX complementation in *L. monocytogenes*. (A) NAD^+^/NADH ratios of indicated *L. monocytogenes* strains +/-exogenous DHNA supplementation grown aerobically at 37°C in defined medium to mid-logarithmic phase. Again, Δ*menB* were spiked with 2 × 10^8^ total CFU from an overnight BHI culture during experimental setup. Data are presented as the standard deviation of the means from three independent experiments. (B) HPLC quantification of fermentation products (Lactate and Acetate) produced and secreted by indicated *L. monocytogenes* strains +/-exogenous DHNA supplementation grown in BHI media aerobically at 37°C to stationary phase. Data are representative of two independent experiments. (C) Indicated *L. monocytogenes* strains (MOI of 10) +/-DHNA supplementation were tested for cytosolic survival in primary IFNAR -/- BMDMs over a 6 hr infection. Data are normalized to wild-type levels of bacteriolysis and presented as the standard deviation of the means from three independent experiments. (D) Intracellular growth of wild-type, Δ*menB*, or Δ*menA* was determined in BMDMs following infection at an MOI of 0.2. Growth curves are representative of at least three independent experiments. Error bars represent the standard deviation of the means of technical triplicates within the representative experiment. ns, not significant.

Having previously observed that DHNA can restore NAD^+^ redox homeostasis and that NOX-dependent NAD^+^ restoration could restore virulence defects of Δ*menB* mutants, we hypothesized that exogenous DHNA supplementation during infection may similarly rescue the cytosolic survival and intracellular growth of DHNA-deficient *L. monocytogenes*. Indeed, the addition of exogenous DHNA during macrophage infection with Δ*menB* or Δ*menI* mutants restored their cytosolic survival back to wild-type and Δ*menA* levels (**Fig. 3C**). Importantly, as observed with NOX expression, DHNA supplementation did not rescue the cytosolic survival of Δ*glmR* mutants whose virulence phenotypes are due to cell wall stress response defects (**Fig. 3C**) (33, 36), demonstrating that the rescue of cytosolic survival by DHNA is specific to DHNA-deficient *L. monocytogenes*. Accordingly, supplementing DHNA during macrophage infection also rescued the ability of Δ*menB* mutants to replicate intracellularly to levels similar of that during Δ*menA* infection (**Fig 3D**). Taken together, these results demonstrate that exogenously provided DHNA can balance NAD^+^/NADH redox homeostasis thereby potentiating *L. monocytogenes* virulence.

### *ndh2* is conditionally essential for DHNA utilization *in vitro*

Although DHNA can drive regeneration of NAD^+^ in *L. monocytogenes* upon exogenous supplementation, it does not restore membrane potential suggesting that it is not simply imported and used to synthesize MK as described in Streptococci (44, 53). We hypothesized that the two annotated *L. monocytogenes*’ NADH dehydrogenases encoded by *ndh1* (*LMRG_02734*) and *ndh2* (*LMRG_02183*), respectively, may utilize DHNA independent of the respiratory pathways to facilitate NAD^+^/NADH homeostasis (23, 25). To test this hypothesis, we generated Δ*ndh1*/*menB*::Tn and *ndh2*::Tn/Δ*menB* mutants and assayed for growth with or without 5μM exogenous DHNA in defined medium. As expected, both double mutants were unable to grow without exogenous DHNA due them being a Δ*menB* mutant (**Fig. 4A**). DHNA supplementation rescued growth of the Δ*ndh1*/*menB*::Tn mutant suggesting that Ndh1 is not required for DHNA-dependent NAD^+^/NADH redox homeostasis. In contrast, the *ndh2*::Tn/Δ*menB* mutant was unable to grow in the presence of exogenous DHNA (**Fig. 4B**). *ndh2* is required for the function of the recently described EET pathway in *L. monocytogenes* (25), therefore we hypothesized that EET may be necessary to utilize DHNA for NAD^+^/NADH redox homeostasis. To test this hypothesis, we transduced *pplA*::Tn, *dmkA*::Tn, *eetA*::Tn, and *fmnA*::Tn mutations into a Δ*menB* background. The growth of all four of these double mutants were rescued upon DHNA supplementation in defined medium (**Fig. S3**).

**Figure 4.**
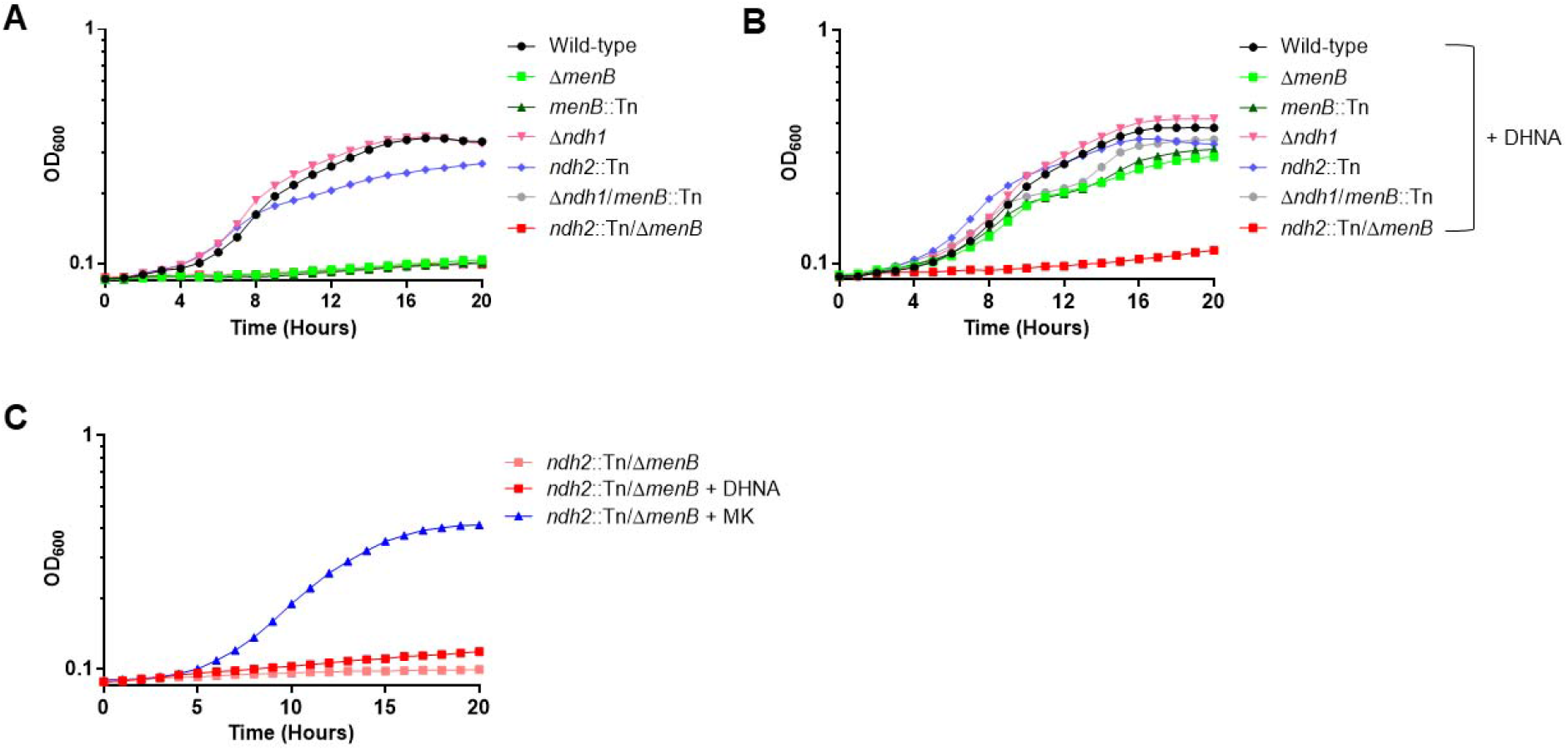
*ndh2* is conditionally essential for DHNA utilization *in vitro*. Indicated strains of *L. monocytogenes* were grown in defined medium without (A) or with (B) 5μM DHNA supplementation aerobically at 37°C and monitored for OD_600_ over 20 hr. (C) *ndh2*::Tn/Δ*menB L. monocytogenes* was grown aerobically in defined medium with either 5μM DHNA or 5μM MK and monitored for growth (OD_600_) over 20 hr. All data represent one representative out of three biological replicates.

Finally, exogenous MK supplementation can restore not only growth of DHNA deficient mutants but also their membrane potential (31, 32), likely through direct insertion of MK in the membrane and subsequent restoration of the aerobic respiratory chain. To ensure that *ndh2*::Tn/Δ*menB* mutants are not more generally incapable of growing in defined media, we supplemented *ndh2*::Tn/Δ*menB* mutants with either DHNA or MK directly.

Supplementation of MK rescued growth of *ndh2*::Tn/Δ*menB* in defined medium unlike DHNA, showing that this mutant is specifically dysfunctional in the use of DHNA as a redox homeostasis substrate (**Fig. 4C**). Taken together, these data suggest that Ndh2 facilitates DHNA-dependent NAD^+^/NADH redox homeostasis in the absence of respiration in *L. monocytogenes*.

## DISCUSSION

Cytosolic pathogens require specific adaptations to survive and replicate within the host. In *L. monocytogenes*, MK biosynthetic intermediate DHNA is among those factors necessary for cytosolic survival, independent of its known role in MK synthesis and cellular respiration (29, 31). In the present study, we sought to address the respiration-independent mechanism by which DHNA is required for the survival and virulence of *L. monocytogenes*. Utilizing a heterologous NOX expression system, we demonstrated that virulence defects associated with loss of DHNA could be rescued by restoration of NAD^+^/NADH homeostasis (**Fig. 1, 2**). We then found that exogenous DHNA supplementation restores NAD^+^/NADH balance, cytosolic survival, and intracellular replication of the DHNA-deficient mutant Δ*menB* (**Fig.3**). Balancing of redox homeostasis also coincided with a marked shift in fermentative flux from lactate to acetate upon DHNA production or supplementation (**Fig. 3B**) to maximize ATP production via SLP through the activity of acetate kinase (34, 37). Lastly, we provide evidence that Ndh2 is the NADH dehydrogenase responsible for restoring redox homeostasis during extracellular DHNA utilization, independent of its role in EET (**Fig. 4**).

Although we’ve demonstrated that Ndh2 is conditionally essential for DHNA utilization in *L. monocytogenes*, it is still unclear how Ndh2 utilizes DHNA to maintain intracellular redox homeostasis. One possibility is that DHNA, or one of its derivatives, may be used as an alternative quinone to directly accept electrons from Ndh2, regenerating NAD^+^ similar to the system recently described in *Shewanella oneidensis* MR-1. Mevers *et al*. recently demonstrated that a derivative of DHNA, 2-amino-3-carboxy-1,4-naphthoquinone (ACNQ), could serve as a novel electron shuttle that functioned to promote redox balance and energy metabolism (38). The authors went on to show that ACNQ is produced non-enzymatically from extracellular DHNA under oxidizing conditions in the presence of a nitrogen donor (i.e. ammonium or amino acids) (38). We have confirmed that indeed, DHNA is secreted by wild-type *L. monocytogenes* (**Fig. S2**) and extracellular DHNA is readily converted to ACNQ in our defined medium based on mass spectrometry analysis (data not shown). Based on this model, it is possible that DHNA produced by *L. monocytogenes* is secreted outside of the cell to shuttle electrons away where it is then freely oxidized non-enzymatically in the local environment to form ACNQ. Newly formed ACNQ would then be imported back into *L. monocytogenes* to be reduced again through the activity of Ndh2. The repeated oxidation and reduction of DHNA and/or ACNQ is the hallmark of an “electron shuttle” and is one of the proposed mechanisms of EET in *S. oneidensis* (38, 39). A strikingly similar model has been described in *Pseudomonas aeruginosa* in which endogenous production of phenazine is cyclically reduced intracellularly, shuttled outside of the cell, and oxidized by a terminal electron acceptor where it is then imported again by the cell (41). Studies to determine whether DHNA/ACNQ fuels an Ndh2-dependent electron shuttle to maintain intracellular redox homeostasis or whether DHNA works via an alternative mechanism are currently ongoing.

It has been proposed that in addition to serving as an electron shuttle by *P. aeruginosa*, secreted phenazine may be used as a shared resource by the surrounding microbial community to fuel their own redox shuttling (42). The function of phenazine as a shared metabolite is also similar to what has been previously documented with the secretion of DHNA being used as a shared resource to fuel metabolic processes of other localized microbes (31, 43–45). Furthermore, a recent study by Tejedor-Sanz *et al*. reported that the homofermentative lactic acid bacteria *Lactiplantibacillus plantarum* contains the EET gene locus previously annotated in *L. monocytogenes*, however it is missing the upstream genes necessary for quinone biosynthesis (40). Upon addition of exogenous DHNA, *L. plantarum* was observed to employ an Ndh2-dependent form of EET that functioned to increase intracellular redox homeostasis by enhancing metabolic flux through fermentative pathways, generating additional lactate, while increasing ATP generation through SLP (40). Importantly, the capacity of DHNA supplementation to induce EET in *L. plantarum* did not coincide with the generation of a PMF to drive oxidative phosphorylation, similar to the phenotypes observed in *L. monocytogenes*. Whether there are functions of *L. monocytogenes* secreted DHNA as a shared metabolite in complex microbial communities such as those found in the intestine during the early stages of infection will require additional future studies.

Overall, we’ve shown that *L. monocytogenes* can utilize DHNA to maintain redox homeostasis through the anaerobic-specific NADH dehydrogenase Ndh2, independent of other EET proteins. Utilization of extracellular DHNA can aid DHNA-deficient *L. monocytogenes* mutants to restore their ability to grow and replicate within the cytosol by potentially driving a yet unclear method of energy metabolism. Pathways involved in unique energy metabolism by various pathogens are increasingly viewed as attractive drug targets and as such future studies utilizing the important model pathogen *L. monocytogenes* to understand the mechanisms of DHNA-dependent redox homeostasis could provide novel insights into the generation of new antimicrobials.

## MATERIALS AND METHODS

### Bacterial strains, plasmid construction, and growth conditions *in vitro*

*L. monocytogenes* strain 10403S is referred to as the wild-type strain, and all other strains used in this study are isogenic derivatives of this parental strain. Vectors were conjugated into *L. monocytogenes* by *Escherichia coli* strain S17 or SM10 (47). The integrative vector pIMK2 was used for constitutive expression of *L. monocytogenes* genes for complementation (48).

*L. monocytogenes* strains were grown at 37°C or 30°C in brain heart infusion (BHI) medium (237500; VWR) or defined medium supplemented with glucose as the sole carbon source. Defined medium is identical to the formulation described by Smith *et al*. (220). *Escherichia coli* strains were grown in Luria-Bertani (LB) broth at 37°C. Antibiotics were used at concentrations of 100 μg/ml carbenicillin (IB02020; IBI Scientific), 10 μg/ml chloramphenicol (190321; MP Biomedicals), 2 μg/ml erythromycin (227330050; Acros Organics), or 30 μg/ml kanamycin (BP906-5; Fisher Scientific) when appropriate. Medium, where indicated, was supplemented with 5 μM 1,4-dihydroxy-2-naphthoate (DHNA) (281255; Sigma) or 5 μM menaquinone (MK) (V9378; Sigma).

### Phage Transduction

Phage transductions were performed as previously described (310). Briefly, MACK *L. monocytogenes* was grown overnight in 3mL LB at 30°C stationary to propagate U153 phage stocks. MACK cultures were pelleted and resuspended in LB + 10mM CaSO_4_ + 10mM MgCl_2_ and added into LB + 0.7% agar + 10mM CaSO_4_ + 10mM MgCl_2_ at 42°C. This mixture was immediately poured on BHI plates and incubated overnight at 30°C. U153 phage plaques were collected and soaked out with 10mM Tris (pH7.5) + 10mM CaSO_4_ + 10mM MgCl_2_. Donor plaque soak-outs were propagated the same way and were filter-sterilized using a 0.2μm syringe filter (09-740-113; Fisher Scientific) and additionally kept sterile by adding 500μL chloroform. Recipient Δ*menB* strain was infected with these donor soak-outs for 30 minutes at room temperature and subsequently plated on BHI agar with erythromycin for selection at 37°C.

### Intracellular bacteriolysis assay

Standard intracellular bacteriolysis assays were performed as previously described (29). Briefly, primary or immortalized bone marrow-derived *IFNAR* ^-/-^ macrophages (5 × 10^5^ per well of 24-well plates) were grown in a monolayer overnight in 500 μL volume. *L. monocytogenes* strains carrying the bacteriolysis reporter pBHE573 (35) were grown at 30°C without shaking overnight. Cultures were then diluted to a final concentration of 5 × 10^8^ CFU/mL in PBS and used to infect macrophages at a MOI of 10. At 1 hr postinfection, media were removed and replaced with media containing 50 μg/ml gentamicin. At 6 hr post infection, media from the wells were aspirated and macrophages were lysed using TNT lysis buffer (20 mM Tris, 200 mM NaCl, 1% Triton [pH 8.0]). Cell lysates were transferred to opaque 96-well plates, and luciferin reagent was added and assayed for luciferase activity (Synergy HT, BioTek; Winooski, VT).

### Intracellular growth assay

Bone marrow-derived macrophages (BMDMs) were prepared from C57BL/6 mice as previously described (51). BMDMs were plated on coverslips at 5 × 10^6^ cells per 60mm dish and allowed to adhere overnight. BMDMs were then infected at an MOI of 0.2 with their respective strain and infection proceeded for 8 hr. At 30 min postinfection, media were removed and replaced with media containing 50 μg/ml gentamicin. Total CFU were quantified at various time points as previously described (50).

### NAD^+^ and NADH measurements

*L. monocytogenes* strains were grown in defined medium at 37°C with shaking to mid-logarithmic phase (OD_600_ 0.4-0.6). Cultures were centrifuged and then resuspended in PBS. Resuspended bacteria were then lysed (2 × 10^8^ total CFU) by a 1:1 addition of 1% dodecyltrimethylammonium bromide (DTAB) (AC409310250; Fisher Scientific) for 5 min with agitation. Lysates were then processed to measure NAD^+^ and NADH levels using the NAD/NADH-Glo assay (Promega, G9071) per the manufacturer’s protocol.

### Fermentation byproduct measurements

Cultures of *L. monocytogenes* were grown in BHI at 37°C with shaking overnight. Bacteria were then centrifuged and 1 mL of the resulting supernatant was filtered through a 0.2μm-pore-size syringe filter (09-740-113; Fisher Scientific). Supernatant samples were next treated with 2μL of H_2_SO_4_ to precipitate any components that might be incompatible with the running buffer. The samples were then centrifuged at 16000 × g for 10 min and then 200μL of each sample transferred to an HPLC vial. HPLC analysis was performed using a ThermoFisher (Waltham, MA) Ultimate 3000 UHPLC system equipped with a UV detector (210 nm). Compounds were separated on a 250 × 4.6 mm Rezex^©^ ROA-Organic acid LC column (Phenomenex Torrance, CA) run with a flow rate of 0.2 mL min^−1^ and at a column temperature of 50 °C. The samples were held at 4 °C prior to injection. Separation was isocratic with a mobile phase of HPLC grade water acidified with 0.015 N H_2_SO_4_ (415 μL L^−1^). At least two standard sets were run along with each sample set. Standards were 100, 20, 4, and 0.8mM concentrations of lactate or acetate. The resultant data was analyzed using the Thermofisher Chromeleon 7 software package.

### Acute virulence assay

All techniques were reviewed and approved by the University of Wisconsin — Madison Institutional Animal Care and Use Committee (IACUC) under the protocol M02501. Female C57BL/6 mice (6 to 8 weeks of age; purchased from Charles River) were used for the purposes of this study. *L. monocytogenes* strains were grown in BHI medium at 30°C without shaking overnight. These cultures were then back-diluted the following day 1:5 into fresh BHI medium and grown at 37°C with shaking until mid-exponential phase (OD_600_ 0.4-0.6). Bacteria were diluted in PBS to a concentration of 5 × 10^5^ CFU/mL and mice were injected intravenously with 1 × 10^5^ total CFU. At 48 hr postinfection, spleens and livers were harvested and homogenized in 0.1% Nonidet P-40 in PBS. Homogenates were then plated on LB plates to enumerate CFU and quantify bacterial burdens.

### Statistical analysis

Statistical significance analysis (GraphPad Prism, version 6.0h) was determined by one-way analysis of variance (ANOVA) with a Dunnett’s posttest comparing wild-type to all other indicated strains or by one-way ANOVA with Tukey’s multiple comparisons test unless otherwise stated (*, P ≤ 0.05; **, P ≤ 0.01; ***, P ≤ 0.001; ****, P ≤ 0.0001).

## ACKNOWLEDGEMENTS

We would like to thank Dr. Samuel Light for providing the vector pPL2-NOX, expressing the water-forming NADH oxidase for integration into *Listeria monocytogenes*.

## FUNDING INFORMATION

This work was funded by the National Institutes of Health (T32007215 [HBS] and R01AI137070 [J-D S]). The funders had no role in study design, data collection and interpretation, or the decision to submit the work for publication.

**Figure S1.**
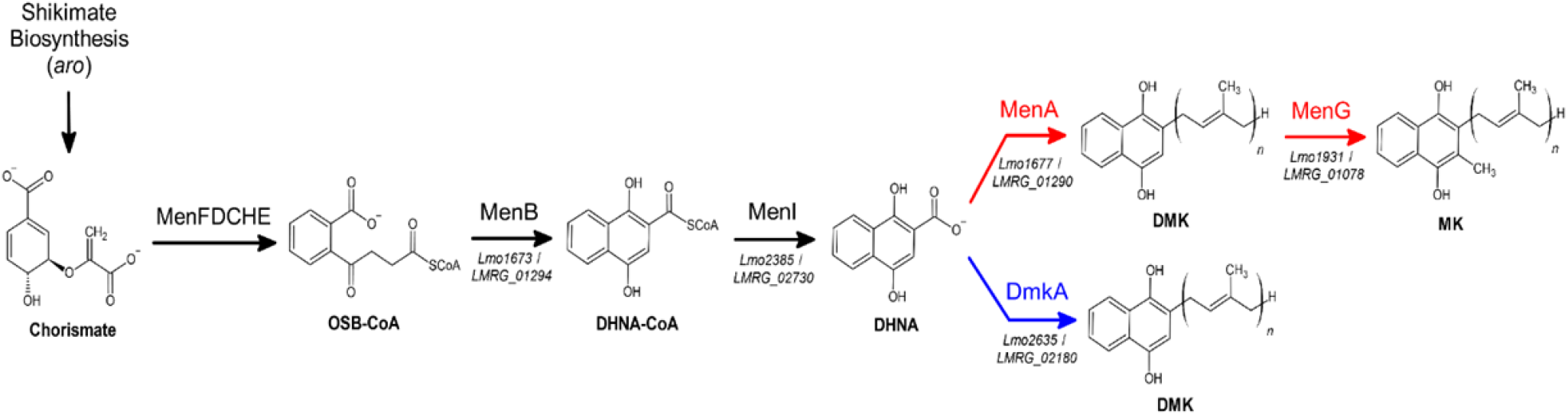
Menaquinone biosynthetic pathway in *Listeria monocytogenes*. Chorismate is generated by the upstream shikimate biosynthesis pathway and is converted to DHNA by the series of listed enzymes (MenFDCHEBI). Red arrows indicate DHNA branching point towards aerobic respiration. Blue arrow indicates DHNA branching point towards anaerobic respiration. Corresponding gene locus numbers for *L. monocytogenes* strains EGD-e (*Lmo*) and 10403S (*LMRG*; parental strain used in this study) are listed underneath reaction arrows. OSB, *o*-succinylbenzoate; DMK, demethylmenaquinone.

**Figure S2.**
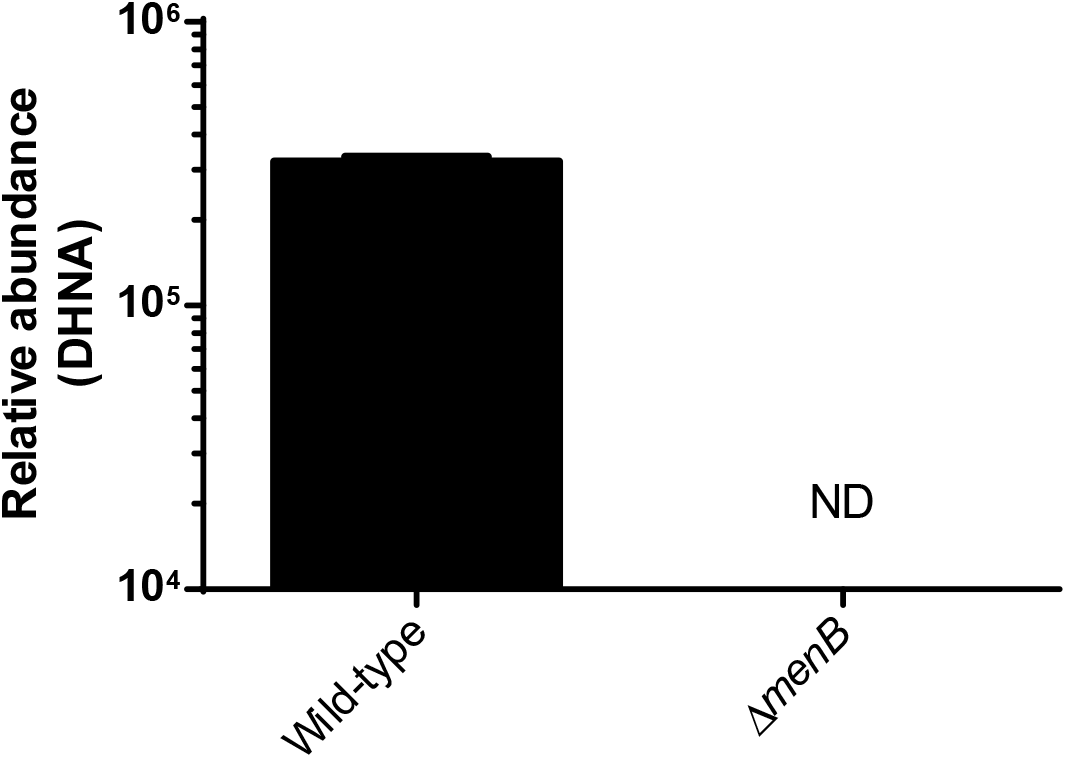
Detection of secreted DHNA by mass spectrometry. Detection of DHNA from the cell-free supernatants of overnight aerobic cultures of wildtype or Δ*menB* strains by mass spectrometry. Data were analyzed via MAVEN. Error bars represent the standard deviation of the means from two independent experiments.

**Figure S3.**
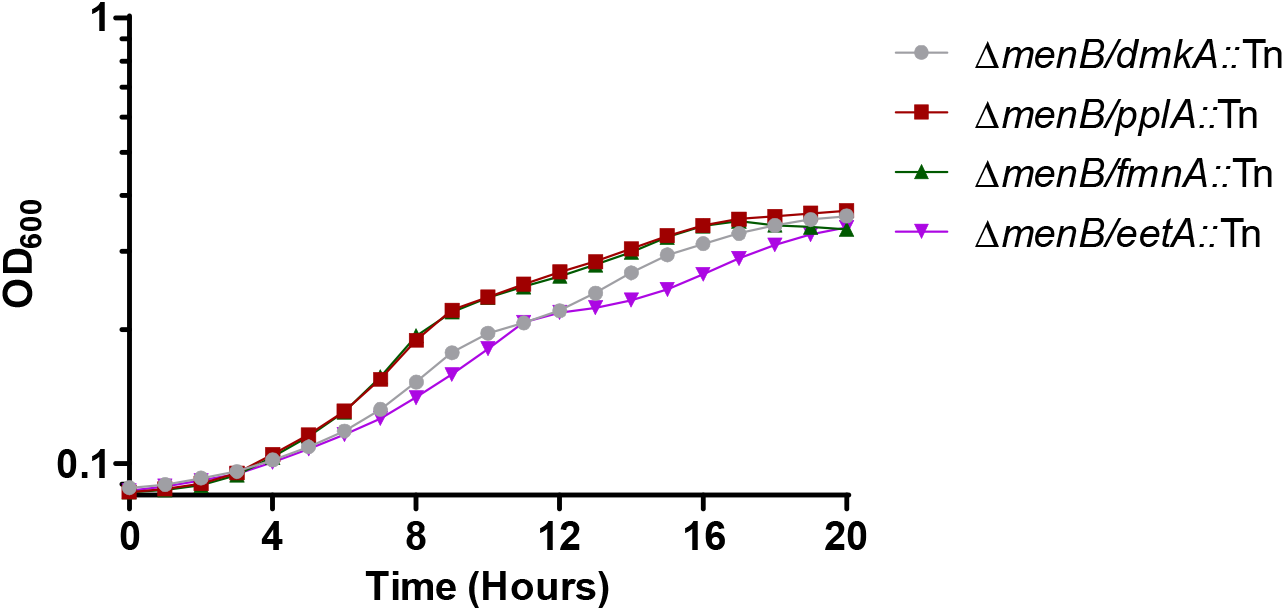
Four separate genes other than *ndh2* that are part of the *L. monocytogenes* EET locus did not display growth defects. Indicated strains were grown in defined medium at 37°C with the addition of 5μM DHNA. OD_600_ was monitored for 20 hours. Data represents one representative out of three biological replicates.

**Table S3.1.** Strains used in this study.

| Strain | Description | Reference |
| --- | --- | --- |
| XL1-Blue | Competent *E. coli* strain | - |
| SM10 | *E. coli* strain for conjugations into *Lm*; Km<sup>R</sup> | (47) |
| 10403S | Parental *L. monocytogenes* (*Lm*) 10403s strain [Wild-type] | - |
| JDS18 | *Lm* with pBHE573 | (35) |
| HS28 | *Lm* with pPL2-NOX | (34) |
| JDS2328 | *Lm* with pBHE573 and pPL2-NOX | (34) |
| JDS25 | *Lm* Δ*glmR* | (35) |
| JDS21 | *Lm* Δ*glmR* with pBHE573 | (35) |
| JDS2327 | *Lm* Δ*glmR* with pPL2-NOX | This work |
| JDS2329 | *Lm* Δ*glmR* with pBHE573 and pPL2-NOX | (34) |
| JDS1161 | *Lm* Δ*menB* | (29) |
| JDS1175 | *Lm* Δ*menB* with pYL116 | (32) |
| JDS1958 | *Lm* Δ*menB* with pPL2-NOX | (34) |
| JDS1191 | *Lm* Δ*menB* with pBHE573 | (29) |
| JDS2333 | *Lm* Δ*menB* with pBHE573 and pPL2-NOX | (34) |
| JDS2240 | *Lm* Δ*menI* | (32) |
| JDS2155 | *Lm* Δ*menI* with pBHE573 | (32) |
| JDS2325 | *Lm* Δ*menI* with pPL2-NOX | This work |
| JDS2326 | *Lm* Δ*menI* with pBHE573 and pPL2-NOX | This work |
| JDS1047 | *Lm* Δ*menA* | (29) |
| JDS813 | *Lm* Δ*menA* with pBHE573 | (29) |
| HS30 | *Lm* Δ*menA* with pPL2-NOX | This work |
| JDS2330 | *Lm* Δ*menA* with pBHE573 and pPL2-NOX | This work |
| JDS1213 | *Lm* Δ*menD* | (52) |
| JDS17 | SM10 *E. coli* with pBHE537 | (35) |
| JDS1957 | SM10 *E. coli* with pPL2-NOX | (34) |
| KL1 | *Lm* Δ*ndh1*/*menB*::Tn | This work |
| KL4 | *Lm* Δ*menB*/*ndh2*::Tn | This work |
| KL9 | *Lm* Δ*menB*/*dmkA*::Tn | This work |
| KL10 | *Lm* Δ*menB*/*pplA*::Tn | This work |
| KL11 | *Lm* Δ*menB*/*fmnA*::Tn | This work |
| KL12 | *Lm* Δ*menB*/*eetA*::Tn | This work |

**Table S3.2.** Plasmids used in this study.

| Plasmids | Description | Reference |
| --- | --- | --- |
| pBHE573 | Bacteriolysis reporter; Cam<sup>R</sup> | (35) |
| pIMK2 | Constitutive expression vector for complementation, P<sub>help</sub>; Kan<sup>R</sup> | (48) |
| pYL116 | *menB* cloned into pIMK2 | (32) |
| pPL2-NOX | NADH oxidase (NOX) cloned into the backbone of pPL2 | (34) |

## References

1. Goetz M, Bubert A, Wang G, Chico-Calero I, Vazquez-Boland JA, Beck M, Slaghuis J, Szalay AA, Goebel W. Microinjection and growth of bacteria in the cytosol of mammalian host cells. Proc Natl Acad Sci U S A. 2001 Oct 9;98(21):12221–6. doi: 10.1073/pnas.211106398.

2. Freitag NE, Port GC, Miner MD. Listeria monocytogenes - from saprophyte to intracellular pathogen. Nat Rev Microbiol. 2009 Sep;7(9):623–8. doi: 10.1038/nrmicro2171.

3. Ray K, Marteyn B, Sansonetti PJ, Tang CM. Life on the inside: the intracellular lifestyle of cytosolic bacteria. Nat Rev Microbiol. 2009 May;7(5):333–40. doi: 10.1038/nrmicro2112.

4. Beuzón CR, Salcedo SP, Holden DW. Growth and killing of a Salmonella enterica serovar Typhimurium sifA mutant strain in the cytosol of different host cell lines. Microbiology. 2002;148. doi:10.1099/00221287-148-9-2705.

5. Brumell JH, Rosenberger CM, Gotto GT, Marcus SL, Finlay BB. SifA permits survival and replication of Salmonella typhimurium in murine macrophages. Cell Microbiol. 2001 Feb;3(2):75–84. doi:10.1046/j.1462-5822.2001.00087.x.

6. Laguna RK, Creasey EA, Li Z, Valtz N, Isberg RR. A Legionella pneumophila-translocated substrate that is required for growth within macrophages and protection from host cell death. Proc Natl Acad Sci U S A. 2006;103. doi:10.1073/pnas.0609012103.

7. Slaghuis J, Goetz M, Engelbrecht F, Goebel W. Inefficient replication of Listeria innocua in the cytosol of mammalian cells. J Infect Dis. 2004 Feb 1;189(3):393–401. doi: 10.1086/381206.

8. Zhang Y, Yeruva L, Marinov A, Prantner D, Wyrick PB, Lupashin V, et al. The DNA Sensor, Cyclic GMP–AMP Synthase, Is Essential for Induction of IFN-β during Chlamydia trachomatis Infection. The Journal of Immunology. 2014;193. doi:10.4049/jimmunol.1302718.

9. Ge J, Gong Y-N, Xu Y, Shao F. Preventing bacterial DNA release and absent in melanoma 2 inflammasome activation by a Legionella effector functioning in membrane trafficking. Proc Natl Acad Sci U S A. 2012;109. doi:10.1073/pnas.1117490109

10. Collins AC, Cai H, Li T, Franco LH, Li X-D, Nair VR, et al. Cyclic GMP-AMP Synthase Is an Innate Immune DNA Sensor for Mycobacterium tuberculosis. Cell Host & Microbe. 2015;17. doi:10.1016/j.chom.2015.05.005

11. Watson RO, Bell SL, MacDuff DA, Kimmey JM, Diner EJ, Olivas J, et al. The Cytosolic Sensor cGAS Detects Mycobacterium tuberculosis DNA to Induce Type I Interferons and Activate Autophagy. Cell Host & Microbe. 2015;17. doi:10.1016/j.chom.2015.05.004

12. Wassermann R, Gulen MF, Sala C, Perin SG, Lou Y, Rybniker J, et al. Mycobacterium tuberculosis Differentially Activates cGAS- and Inflammasome-Dependent Intracellular Immune Responses through ESX-1. Cell Host & Microbe. 2015;17. doi:10.1016/j.chom.2015.05.003

13. McNab, F., Mayer-Barber, K., Sher, A. et al. Type I interferons in infectious disease. Nat Rev Immunol 15, 87–103 (2015). https://doi.org/10.1038/nri3787

14. Portnoy DA, Auerbuch V, Glomski IJ. The cell biology of Listeria monocytogenes infection: the intersection of bacterial pathogenesis and cell-mediated immunity. J Cell Biol. 2002 Aug 5;158(3):409–14. doi: 10.1083/jcb.200205009.

15. Vázquez-Boland JA, Kuhn M, Berche P, Chakraborty T, Domínguez-Bernal G, Goebel W, González-Zorn B, Wehland J, Kreft J. Listeria pathogenesis and molecular virulence determinants. Clin Microbiol Rev. 2001 Jul;14(3):584–640. doi: 10.1128/CMR.14.3.584-640.2001.

16. Portnoy DA, Jacks PS, Hinrichs DJ. Role of hemolysin for the intracellular growth of Listeria monocytogenes. J Exp Med. 1988 Apr 1;167(4):1459–71. doi: 10.1084/jem.167.4.1459.

17. Tilney LG, Portnoy DA. Actin filaments and the growth, movement, and spread of the intracellular bacterial parasite, Listeria monocytogenes. J Cell Biol. 1989 Oct;109(4 Pt 1):1597–608. doi: 10.1083/jcb.109.4.1597.

18. Brundage RA, Smith GA, Camilli A, Theriot JA, Portnoy DA. Expression and phosphorylation of the Listeria monocytogenes ActA protein in mammalian cells. Proc Natl Acad Sci U S A. 1993 Dec 15;90(24):11890–4. doi: 10.1073/pnas.90.24.11890.

19. Moors MA, Levitt B, Youngman P, Portnoy DA. Expression of listeriolysin O and ActA by intracellular and extracellular Listeria monocytogenes. Infect Immun. 1999 Jan;67(1):131–9. doi: 10.1128/IAI.67.1.131-139.1999.

20. Shetron-Rama LM, Marquis H, Bouwer HG, Freitag NE. Intracellular induction of Listeria monocytogenes actA expression. Infect Immun. 2002 Mar;70(3):1087–96. doi: 10.1128/IAI.70.3.1087-1096.2002.

21. Smith GA, Marquis H, Jones S, Johnston NC, Portnoy DA, Goldfine H. The two distinct phospholipases C of Listeria monocytogenes have overlapping roles in escape from a vacuole and cell-to-cell spread. Infect Immun. 1995 Nov;63(11):4231–7. doi: 10.1128/iai.63.11.4231-4237.1995.

22. Sauer JD, Herskovits AA, O’Riordan MXD. Metabolism of the Gram-Positive Bacterial Pathogen Listeria monocytogenes. Microbiol Spectr. 2019 Jul;7(4):10.1128/microbiolspec.GPP3-0066-2019. doi: 10.1128/microbiolspec.GPP3-0066-2019.

23. P G, L F, C B, C R, A A, F B, P B, H B, P B, T C, A C, F C, E C, A de D, P D, E D, G D-B, E D, L D, O D, KD E, H F, F GP, P G, L G, W G, N G-L, T H, J H, D J, LM J, U K, J K, M K, F K, G K, E M, A M, JM V, E N, H N, G N, S N, B de P, JC P-D, R P, B R, M R, T S, N S, A T, JA V-B, H V, J W, P C. 2001. Comparative genomics of Listeria species. Science. 2001 Oct 26;294(5543):849–52. doi: 10.1126/science.1063447.

24. Romick TL, Fleming HP, McFeeters RF. Aerobic and anaerobic metabolism of Listeria monocytogenes in defined glucose medium. Appl Environ Microbiol. 1996 Jan;62(1):304–7. doi: 10.1128/aem.62.1.304-307.1996.

25. Light SH, Su L, Rivera-Lugo R, Cornejo JA, Louie A, Iavarone AT, Ajo-Franklin CM, Portnoy DA. A flavin-based extracellular electron transfer mechanism in diverse Gram-positive bacteria. Nature. 2018 Oct;562(7725):140–144. doi: 10.1038/s41586-018-0498-z.

26. Meganathan R. Biosynthesis of menaquinone (vitamin K2) and ubiquinone (coenzyme Q): a perspective on enzymatic mechanisms. Vitam Horm. 2001;61:173–218. doi: 10.1016/s0083-6729(01)61006-9.

27. Corbett D, Goldrick M, Fernandes VE, Davidge K, Poole RK, Andrew PW, Cavet J, Roberts IS. Listeria monocytogenes Has Both Cytochrome bd-Type and Cytochrome aa3-Type Terminal Oxidases, Which Allow Growth at Different Oxygen Levels, and Both Are Important in Infection. Infect Immun. 2017 Oct 18;85(11):e00354–17. doi: 10.1128/IAI.00354-17.

28. Light SH, Méheust R, Ferrell JL, Cho J, Deng D, Agostoni M, Iavarone AT, Banfield JF, D’Orazio SEF, Portnoy DA. 2019. Extracellular electron transfer powers flavinylated extracellular reductases in Gram-positive bacteria. Proc Natl Acad Sci U S A 116:26892–26899.

29. Chen, G.Y., McDougal, C.E., D’Antonio, M.A., Portman, J.L. and Sauer, JD. A Genetic Screen Reveals that Synthesis of 1,4-Dihydroxy-2-Naphthoate (DHNA), but Not Full-Length Menaquinone, Is Required for Listeria monocytogenes Cytosolic Survival. MBio. 2017 Mar 21;8(2).

30. Stritzker J, Janda J, Schoen C, Taupp M, Pilgrim S, Gentschev I, Schreier P, Geginat G, Goebel W. Growth, virulence, and immunogenicity of Listeria monocytogenes aro mutants. Infect Immun. 2004 Oct;72(10):5622–9. doi: 10.1128/IAI.72.10.5622-5629.2004.

31. Chen GY, Kao CY, Smith HB, Rust DP, Powers ZM, Li AY, Sauer JD. Mutation of the Transcriptional Regulator YtoI Rescues Listeria monocytogenes Mutants Deficient in the Essential Shared Metabolite 1,4-Dihydroxy-2-Naphthoate (DHNA). Infect Immun. 2019 Dec 17;88(1):e00366–19. doi: 10.1128/IAI.00366-19.

32. Smith HB, Li TL, Liao MK, Chen GY, Guo Z, Sauer JD. Listeria monocytogenes MenI Encodes a DHNA-CoA Thioesterase Necessary for Menaquinone Biosynthesis, Cytosolic Survival, and Virulence. Infect Immun. 2021 Apr 16;89(5):e00792–20. doi: 10.1128/IAI.00792-20.

33. Pensinger DA, Gutierrez K V., Smith HB, Vincent WJB, Stevenson DS, Black KA, Perez-Medina KM, Dillard JP, Rhee KY, Amador-Noguez D, Huynh TN, Sauer J-D. 2021. Listeria monocytogenes GlmR is an accessory uridyltransferase essential for cytosolic survival and virulence. bioRxiv. https://doi.org/10.1101/2021.10.27.466214.

34. Rivera-Lugo R, Deng D, Anaya-Sanchez A, Tejedor-Sanz S, Tang E, Reyes Ruiz VM, Smith HB, Titov DV, Sauer JD, Skaar EP, Ajo-Franklin CM, Portnoy DA, Light SH. Listeria monocytogenes requires cellular respiration for NAD^+^ regeneration and pathogenesis. Elife. 2022 Apr 5;11:e75424. doi: 10.7554/eLife.75424.

35. Sauer JD, Witte CE, Zemansky J, Hanson B, Lauer P, Portnoy DA. Listeria monocytogenes triggers AIM2-mediated pyroptosis upon infrequent bacteriolysis in the macrophage cytosol. Cell Host Microbe. 2010 May 20;7(5):412–9. doi: 10.1016/j.chom.2010.04.004.

36. Pensinger DA, Boldon KM, Chen GY, Vincent WJ, Sherman K, Xiong M, Schaenzer AJ, Forster ER, Coers J, Striker R, Sauer JD. The Listeria monocytogenes PASTA Kinase PrkA and Its Substrate YvcK Are Required for Cell Wall Homeostasis, Metabolism, and Virulence. PLoS Pathog. 2016 Nov 2;12(11):e1006001. doi: 10.1371/journal.ppat.1006001.

37. Hunt KA, Flynn JM, Naranjo B, Shikhare ID, Gralnick JA. Substrate-level phosphorylation is the primary source of energy conservation during anaerobic respiration of Shewanella oneidensis strain MR-1. J Bacteriol. 2010 Jul;192(13):3345–51. doi: 10.1128/JB.00090-10. Epub 2010 Apr 16.

38. Mevers E, Su L, Pishchany G, Baruch M, Cornejo J, Hobert E, Dimise E, Ajo-Franklin CM, Clardy J. An elusive electron shuttle from a facultative anaerobe. Elife. 2019 Jun 24;8:e48054. doi: 10.7554/eLife.48054.

39. Brutinel ED, Gralnick JA. Shuttling happens: soluble flavin mediators of extracellular electron transfer in Shewanella. Appl Microbiol Biotechnol. 2012 Jan;93(1):41–8. doi: 10.1007/s00253-011-3653-0.

40. Tejedor-Sanz S, Stevens ET, Li S, Finnegan P, Nelson J, Knoesen A, Light SH, Ajo-Franklin CM, Marco ML. Extracellular electron transfer increases fermentation in lactic acid bacteria via a hybrid metabolism. Elife. 2022 Feb 11;11:e70684. doi: 10.7554/eLife.70684.

41. Wang Y, Kern SE, Newman DK. Endogenous phenazine antibiotics promote anaerobic survival of Pseudomonas aeruginosa via extracellular electron transfer. J Bacteriol. 2010 Jan;192(1):365–9. doi: 10.1128/JB.01188-09.

42. Rabaey K, Boon N, Höfte M, Verstraete W. Microbial phenazine production enhances electron transfer in biofuel cells. Environ Sci Technol. 2005 May 1;39(9):3401–8. doi: 10.1021/es048563o.

43. Isawa K, Hojo K, Yoda N, Kamiyama T, Makino S, Saito M, Sugano H, Mizoguchi C, Kurama S, Shibasaki M, Endo N, Sato Y. Isolation and identification of a new bifidogenic growth stimulator produced by Propionibacterium freudenreichii ET-3. Biosci Biotechnol Biochem. 2002 Mar;66(3):679–81. doi: 10.1271/bbb.66.679.

44. Franza T, Delavenne E, Derré-Bobillot A, Juillard V, Boulay M, Demey E, Vinh J, Lamberet G, Gaudu P. A partial metabolic pathway enables group b streptococcus to overcome quinone deficiency in a host bacterial community. Mol Microbiol. 2016 Oct;102(1):81–91. doi: 10.1111/mmi.13447.

45. Kang JE, Kim TJ, Moon GS. A Novel Lactobacillus casei LP1 Producing 1,4-Dihydroxy-2-Naphthoic Acid, a Bifidogenic Growth Stimulator. Prev Nutr Food Sci. 2015 Mar;20(1):78–81. doi: 10.3746/pnf.2015.20.1.78.

46. Grubmüller S, Schauer K, Goebel W, Fuchs TM, Eisenreich W. Analysis of carbon substrates used by Listeria monocytogenes during growth in J774A.1 macrophages suggests a bipartite intracellular metabolism. Front Cell Infect Microbiol. 2014 Nov 3;4:156. doi: 10.3389/fcimb.2014.00156.

47. Simon, R., Priefer, U. and Pühler, A. A Broad Host Range Mobilization System for in Vivo Genetic Engineering: Transposon Mutagenesis in Gram Negative Bacteria. Nature Biotechnology, 1983, 1, 784–791. http://dx.doi.org/10.1038/nbt1183-784

48. Monk IR, Gahan CG, Hill C. Tools for functional postgenomic analysis of listeria monocytogenes. Appl Environ Microbiol. 2008 Jul;74(13):3921–34. doi: 10.1128/AEM.00314-08. Epub 2008 Apr 25.

49. Hodgson DA. Generalized transduction of serotype 1/2 and serotype 4b strains of Listeria monocytogenes. Mol Microbiol. 2000 Jan;35(2):312–23. doi: 10.1046/j.1365-2958.2000.01643.x.

50. Rohmer L, Hocquet D, Miller SI. Are pathogenic bacteria just looking for food? Metabolism and microbial pathogenesis. Trends Microbiol. 2011 Jul;19(7):341–8. doi: 10.1016/j.tim.2011.04.003. Epub 2011 May 18.

51. Jones S, Portnoy DA. Characterization of Listeria monocytogenes pathogenesis in a strain expressing perfringolysin O in place of listeriolysin O. Infect Immun. 1994 Dec;62(12):5608–13. doi: 10.1128/iai.62.12.5608-5613.1994.

52. Perry KJ, Higgins DE. A differential fluorescence-based genetic screen identifies Listeria monocytogenes determinants required for intracellular replication. J Bacteriol. 2013 Aug;195(15):3331–40. doi: 10.1128/JB.00210-13.

53. Lencina AM, Franza T, Sullivan MJ, Ulett GC, Ipe DS, Gaudu P, Gennis RB, Schurig-Briccio LA. Type 2 NADH Dehydrogenase Is the Only Point of Entry for Electrons into the Streptococcus agalactiae Respiratory Chain and Is a Potential Drug Target. mBio. 2018 Jul 3;9(4):e01034–18. doi: 10.1128/mBio.01034-18.

